# Bespoke conformation and antibody recognition distinguishes the streptococcal immune evasion factors EndoS and EndoS2

**DOI:** 10.1101/2023.08.15.553389

**Authors:** Abigail S. L. Sudol, Ivo Tews, Max Crispin

**Author notes:** Correspondence to Ivo Tews and Max Crispin.

## Abstract

The IgG-specific endoglycosidases EndoS and EndoS2 from *Streptococcus pyogenes* ablate IgG function by removing the conserved N-linked glycans present on the Fc region. Their role in immune evasion, by inactivation of IgG antibodies, has led these enzymes to be investigated as therapeutics for suppressing unwanted immune activation. Their activity and precise substrate specificity has also prompted the development of these enzymes as tools for engineering IgG glycosylation. Recent structural studies have revealed how EndoS drives specificity for IgG by binding the Fc peptide surface with a domain that has homology for a carbohydrate-binding module (CBM). Here, we present the crystal structure of the EndoS2-IgG1 Fc complex at 3.0 Å resolution. Comparison with the EndoS-IgG1 Fc structure reveals a similar mode of interaction, but slightly different orientations resulting from different interfaces with glycosidase and CBM domains, leading to recognition of distinct Fc surfaces. These findings rationalise previous observations that non-catalytic domains cannot readily be substituted. The structural information presented here will guide the continued development of IgG-specific endoglycosidases in antibody glycoengineering and immunotherapy.

## Main text

The pathogen *Streptococcus pyogenes* is highly adapted for human infection and has thus developed a range of immune evasion mechanisms for prolonging infection [1]. This pathogenic bacterial species possesses a toolkit of enzymes which specifically target and degrade immunoglobulin G (IgG) antibodies, the most abundant antibody found within human serum [2]. For example, the protease IgG-degrading enzyme of *S. pyogenes* (IdeS) deactivates IgG by cleaving within the lower hinge region, yielding F(ab′)_2_ and Fc fragments [3, 4]. Another protease, SpeB, also cleaves a broader range of immunoglobulins and other components of the immune system [5, 6].

Streptococcal EndoS [7] and EndoS2 [8] enzymes are additionally utilised by the bacterium for the removal of N-linked glycans from IgG Fc. Specifically, these enzymes cleave the β1−4 glycosidic linkage between the two *N*-acetylglucosamine (GlcNAc) residues within the glycan core, releasing the majority of the glycan and leaving a single GlcNAc, variably modified with fucose. EndoS is specific towards complex-type, biantennary glycans on IgG Fc [7, 9], whereas EndoS2 exhibits broader glycan specificity, also cleaving other classes of N-linked glycans such as oligomannose-type and hybrid-type [8, 10]. The broader substrate specificity exhibited by EndoS2 appears to be possible due to a larger glycan-binding groove within the active site which can accommodate such bulkier glycans [11], in comparison to that of EndoS [12]. The glycosylation of IgG occurs at residue N297 within the Fc Cγ2 domains and is conserved across all IgG subtypes [13]. Glycosylation of the Fc domain is implicated in maintaining Fc structural integrity [14-19], Fc gamma receptor (FcγR)-and complement-mediated immune activation [2, 17, 20]. Removal of the glycan therefore impedes Fc-mediated effector functions [19, 21], although activity can sometimes be detected for particular IgG subclasses when deglycosylated Fcs are presented in immune complexes [22].

EndoS was demonstrated to improve survival of *S. pyogenes* in an opsonophagocytic assay, due to reduced FcγR-and complement-mediated immune activation [23]. Moreover, IgG glycan hydrolysis by EndoS has been shown to promote virulence and survival of the bacterium *in vivo* [24], thus demonstrating its role in immune evasion. This highlights the potential of EndoS, and EndoS2, as therapeutic agents for dampening unwanted immune activation, such as during organ transplantation or in an autoimmune disease context. Indeed, EndoS has been successfully utilised as a treatment in several pre-clinical models of autoimmune disease [25-34], and, in combination with IdeS, was shown to be successful at inactivating donor-specific antibodies in a murine model of bone marrow transplantation [35]. The specific deactivation of competing serum IgG by EndoS and/or IdeS is also being investigated for the potentiation of therapeutic antibodies [19, 36].

Additional interest in these enzymes stems from potential use in biotechnological application, providing tools for antibody glycoengineering [37, 38]. Wild-type enzymes can be used to cleave off unwanted glycoforms, and transglycosylation reactions can be performed on this antibody scaffold to generate desired glycoforms [39-43]. Such transglycosylation reactions have been optimised using variants of these enzymes, although both wild-type EndoS [9, 44] and EndoS2 [41] have been shown to possess some transglycosylation activity. A further development has been the use of wild-type EndoS2 in “one-step” reactions for synthesis of antibody-drug conjugates [45, 46]. The precise control of antibody glycosylation has been applied in several clinically-used antibodies for improved immune effector function [47-51], thus demonstrating their utility in antibody engineering.

The potential therapeutic and biotechnological uses of EndoS and EndoS2 have prompted extensive research into the structure and function of these enzymes. These endoglycosidases are multi-domain enzymes, with the catalytic glycosidase domain and the so-called carbohydrate binding module (CBM) being of particular interest [12, 52-54]. Recent structural studies revealed that the functional role of the CBM in EndoS is not to bind carbohydrate, but rather to specify peptide binding on the Fc surface [55, 56], which provided a structural rationale for the abrogated enzymatic activity of EndoS lacking the CBM [52, 53], further explaining the inability of EndoS to cleave glycans from denatured IgG [6]. Similarly, it is known for that the CBM of EndoS2 is essential for activity. Furthermore, the CBM must work in tandem with the catalytic domain, as a chimeric enzyme displacing the EndoS2 catalytic domain onto the EndoS scaffold leads to a non-functional enzyme [11].

We therefore sought to investigate the structural basis behind the specificity of EndoS2 for IgG, and how it differs from EndoS in substrate recognition. We used the previously established “less-crystallisable” Fc to obtain crystals of the EndoS2-IgG1 Fc complex [55]. We further introduced a L234C exchange, in an attempt stabilise the flexible hinge region through disulphide formation (see Supplementary Information). The enzyme EndoS2_38–843_was expressed as an inactive exchange variant D184A/E186L with C-terminal His-tag (see SI, [11]). Purified EndoS2^D184A/E186L^ and IgG1 Fc^L234C/E382A^ were combined in a 2:1 molar ratio and the resulting complex purified by size exclusion chromatography and submitted to crystallisation for structure determination (see SI).

Crystallographic studies of enzymes in complex with their glycan substrate have provided insight into the mechanism of action of endoglycosidases [11, 12, 57-59]. Our recent structure of EndoS in complex with its IgG1 Fc substrate [55] and a cryo-electron microscopy study [56] provided further insight. Here, we present the crystal structure of EndoS2 in complex with IgG1 Fc substrate at a resolution of 3.0 Å (Table 1), with N-linked glycans modelled using the carbohydrate module in Coot [60] and validated using Privateer [61], as shown in Figure 1. The complex crystallised in space group *P*4_3_2_1_2, with three copies of the EndoS2-half Fc complex within the asymmetric unit (Table 1).

**Table 1:**
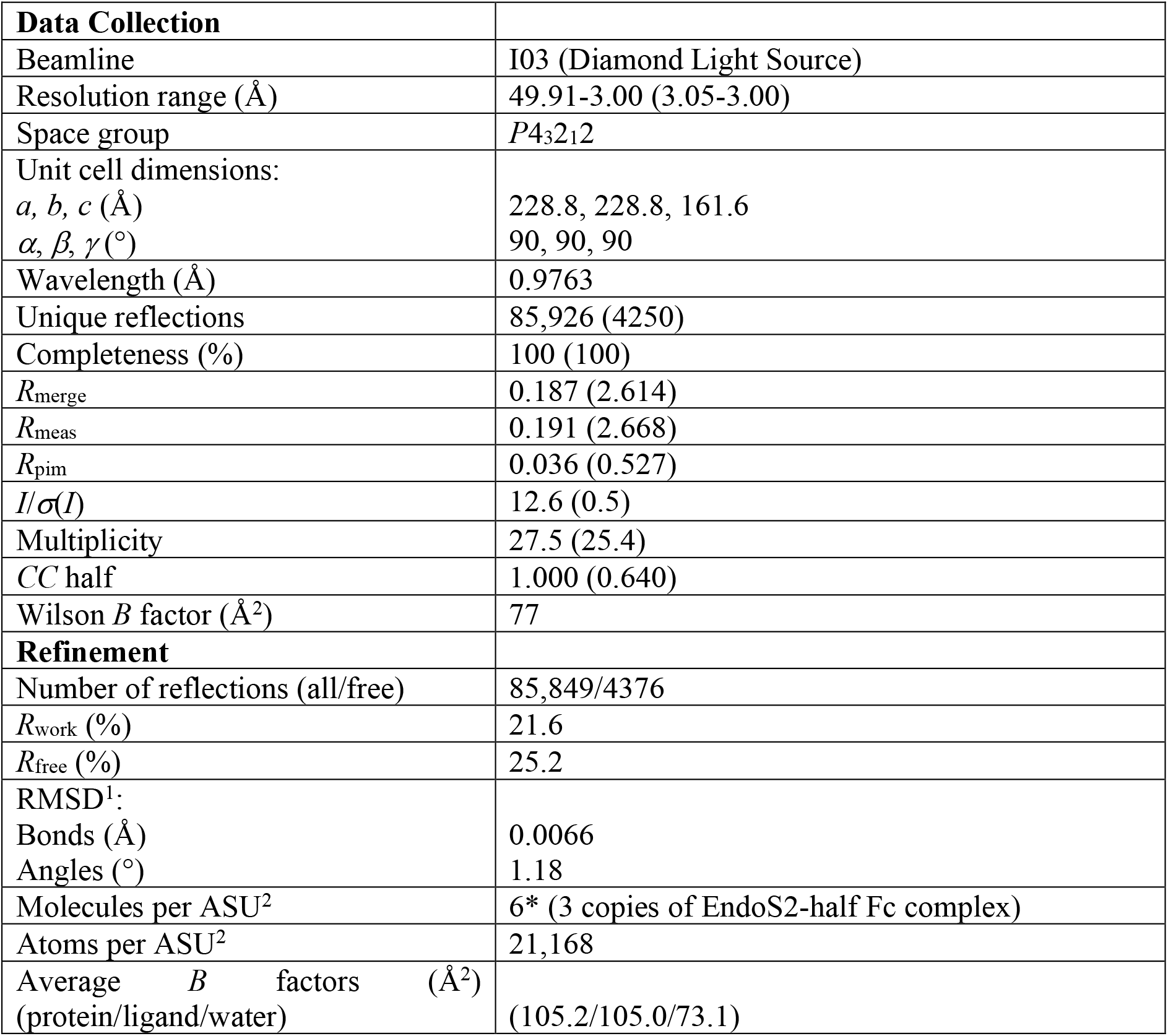
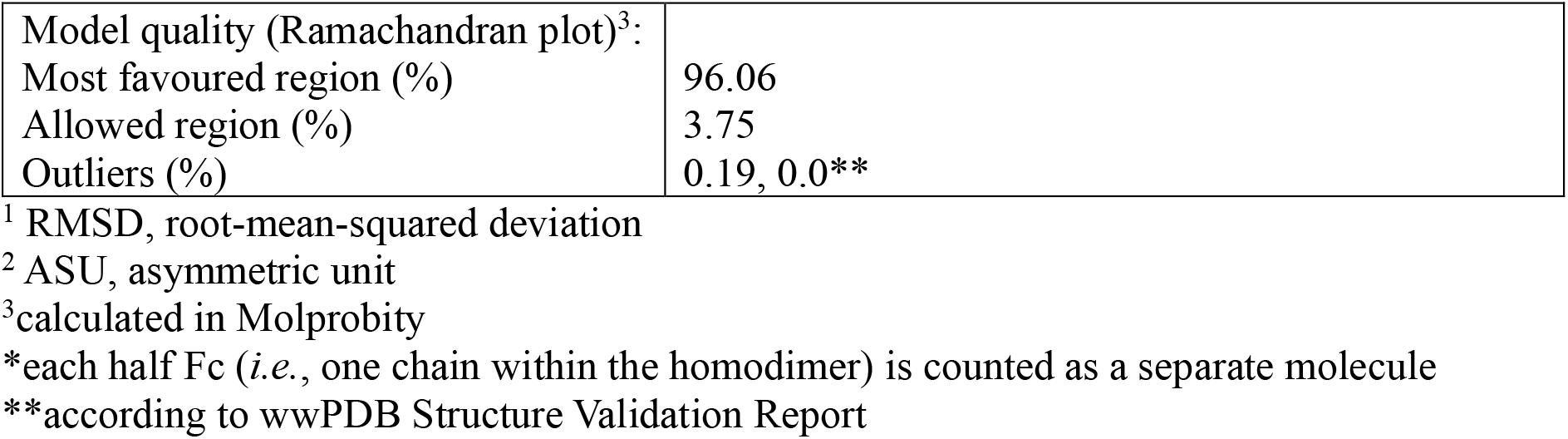
Crystallographic data collection and refinement statistics for EndoS2^D184A/E186L^-IgG1 Fc^L234C/E382A^ complex. Values for the highest resolution shell are reported in parentheses.

**Figure 1:**
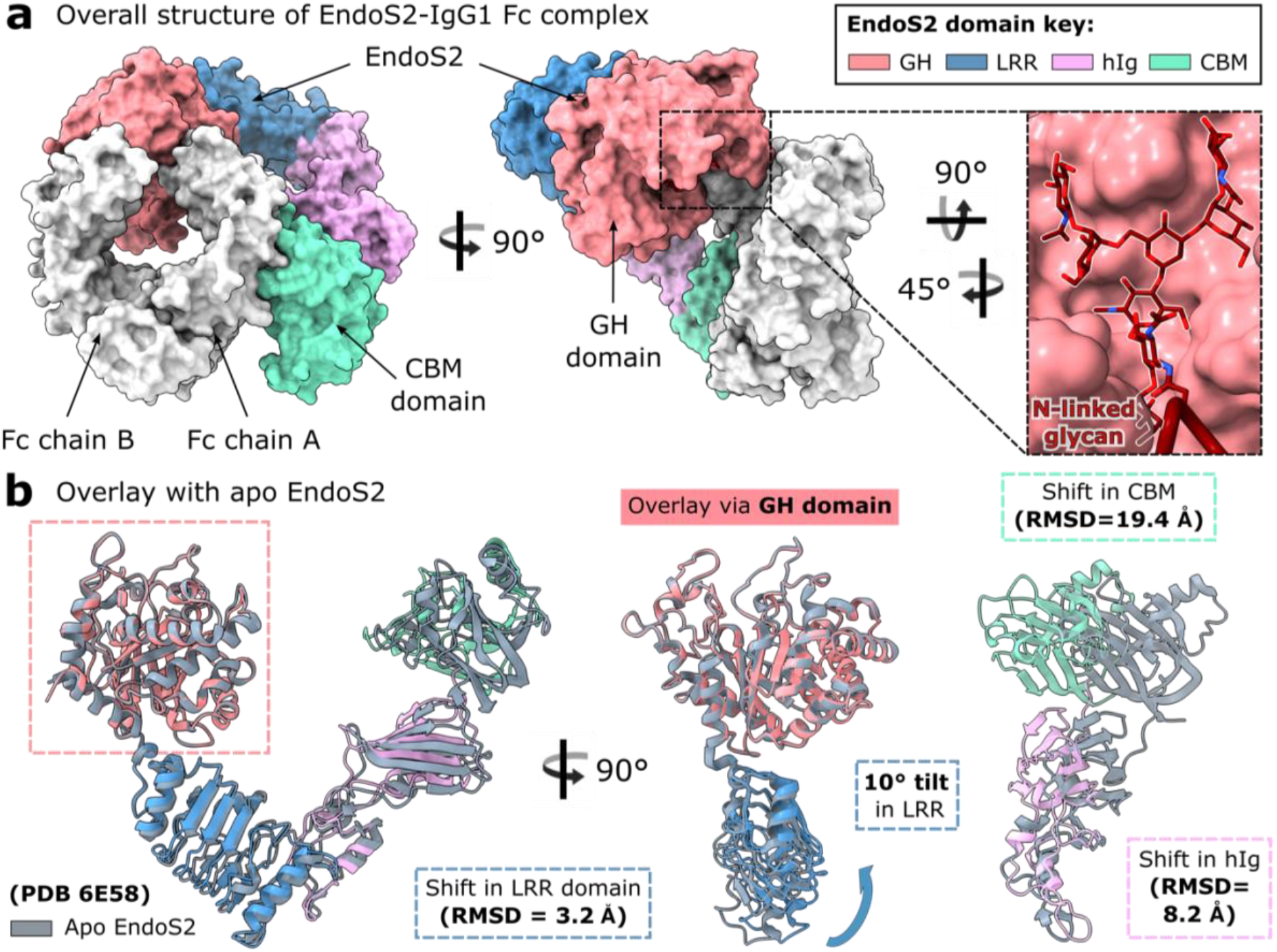
Overall structure of EndoS2^D184A/E186L^-IgG1 Fc^L234C/E382A^ complex. **a** Surface representation of the EndoS-IgG1 Fc complex. Two EndoS2 molecules bind each Fc homodimer in the crystal (see Supplementary Figure 2); for clarity, only one EndoS2 molecule is shown here. IgG1 Fc is depicted in silver, with the N297 glycan depicted as maroon sticks and coloured by heteroatom (oxygen, red; nitrogen, blue). EndoS2 domains are coloured as follows: glycoside hydrolase (GH), coral; leucine-rich repeat (LRR), blue; hybrid immunoglobulin (hIg), lilac; carbohydrate-binding module (CBM), green. **b** Superposition of apo EndoS2 (PDB 6E58) and EndoS2 in complex with IgG1 Fc, with respect to the GH domain (aligned Cαs for EndoS2 residues 46*–*386). IgG1 Fc in complexed EndoS2 is omitted here for clarity. The figure reveals the shift of GH and CBM relative to each other, with the entire GH-LRR-hIg-CBM scaffold in different orientation. RMSDs and plane angle differences for EndoS2 domain shifts after these superpositions are indicated, as calculated in UCSF ChimeraX. Images depicting protein structure were prepared using UCSF ChimeraX [62].

EndoS2 comprises a glycosylhydrolase domain (GH, coral, residues 43–386), a leucine-rich repeat domain (LRR, blue, residues 387–547), a hybrid immunoglobulin domain (hIg, lilac, residues 548–680) and a carbohydrate binding module (CBM, green, residues 681–843) [11]. The enzyme is shaped like a “V”, and captures one Cγ2 domain of the Fc through binding by the GH and CBM domains (Figure 1), similar to that observed in the structure of the EndoS-Fc complex [55, 56]. Each Fc γ-chain is seen to interact with one EndoS2 molecule, similar to what was observed with the EndoS-Fc crystal lattice [55].

In the EndoS2-IgG1 Fc complex, the uncleaved Fc N-linked glycan is captured in a “flipped-out” conformation and bound within the GH domain groove (Figure 1a), which was identified by co-crystals of EndoS2 in complex with glycan substrates [11]. This observation is in stark contrast to the typically-observed position of Fc glycans within crystal structures, in which they appear to sit in-between the Cγ2 domains and interact with Fc surface residues [13, 63-65]. The carbohydrate is well-ordered, displaying low *B* factors with respect to the average *B* factor of the complex (105.8 Å^2^, Table 1). A polder map additionally displays clear density consistent with an uncleaved glycan in this conformation (Supplementary Figure 3). Overall, this demonstrates the same mode of glycan capture as observed in the EndoS-IgG1 Fc complex [55], and further corroborates literature reporting heterogeneity of Fc N-linked glycan conformations [66-70].

Superposition of complexed and apo EndoS2 (PDB 6E58) reveals a shifting of multiple domains upon Fc binding (Figure 1b). An overlay with respect to the GH domain (calculated by aligning Cαs for residues 46–386) results in a 10° tilt of the LRR (with respect to its position in the apo structure). Consequently, the hIg and CBM are also displaced (with RMSDs of 8.2 Å and 19.4 Å, respectively). An overlay with respect to the CBM (calculated by aligning Cαs for residues 681–843) results in smaller domain shifts: the adjacent hIg domain is tilted by 4.2°, while the LRR and GH are displaced by 5.1 Å and 6.3 Å, respectively. Therefore, both the GH and CBM domains are rearranged with respect to the rest of the enzyme upon Fc binding. The conformation of the LRR-hIg scaffold is also affected: an overlay with respect to the LRR domain (calculated by aligning Cαs for residues 387–547) results in a 11.6° tilt in the hIg. Thus, the relative domain positioning within EndoS2 is shifted upon binding the Fc substrate, although discrete domains can be fully superimposed and so do not undergo individual conformational changes. Analysis of apo and glycan-bound EndoS2 crystal structures has revealed no conformational changes upon glycan binding [11], which indicates that EndoS2 domain shifts observed here are solely due to binding the Fc peptide surface.

The binding interface of the EndoS2-IgG1 Fc complex is analysed in Figure 2. The GH and CBM domains within EndoS2 both contact the same Fc Cγ2 domain. This corroborates hydrogen-deuterium exchange data of the EndoS2-rituximab complex, which suggested a role for both domains in complex formation [11]. Crystal structures of EndoS2 in complex with oligomannose-and complex-type glycan substrates have identified GH amino acids involved in glycan binding [11]; the EndoS2-Fc complex structure additionally reveals interactions with the first GlcNAc and fucose moieties within the glycan (Figure 2b). The aromatic side chains of F189, Y251, Y252 and W297 provide hydrophobic stacking interactions, while the Y252 backbone and Q255 side chain form hydrogen bonds with the fucose (Figure 2b).

**Figure 2:**
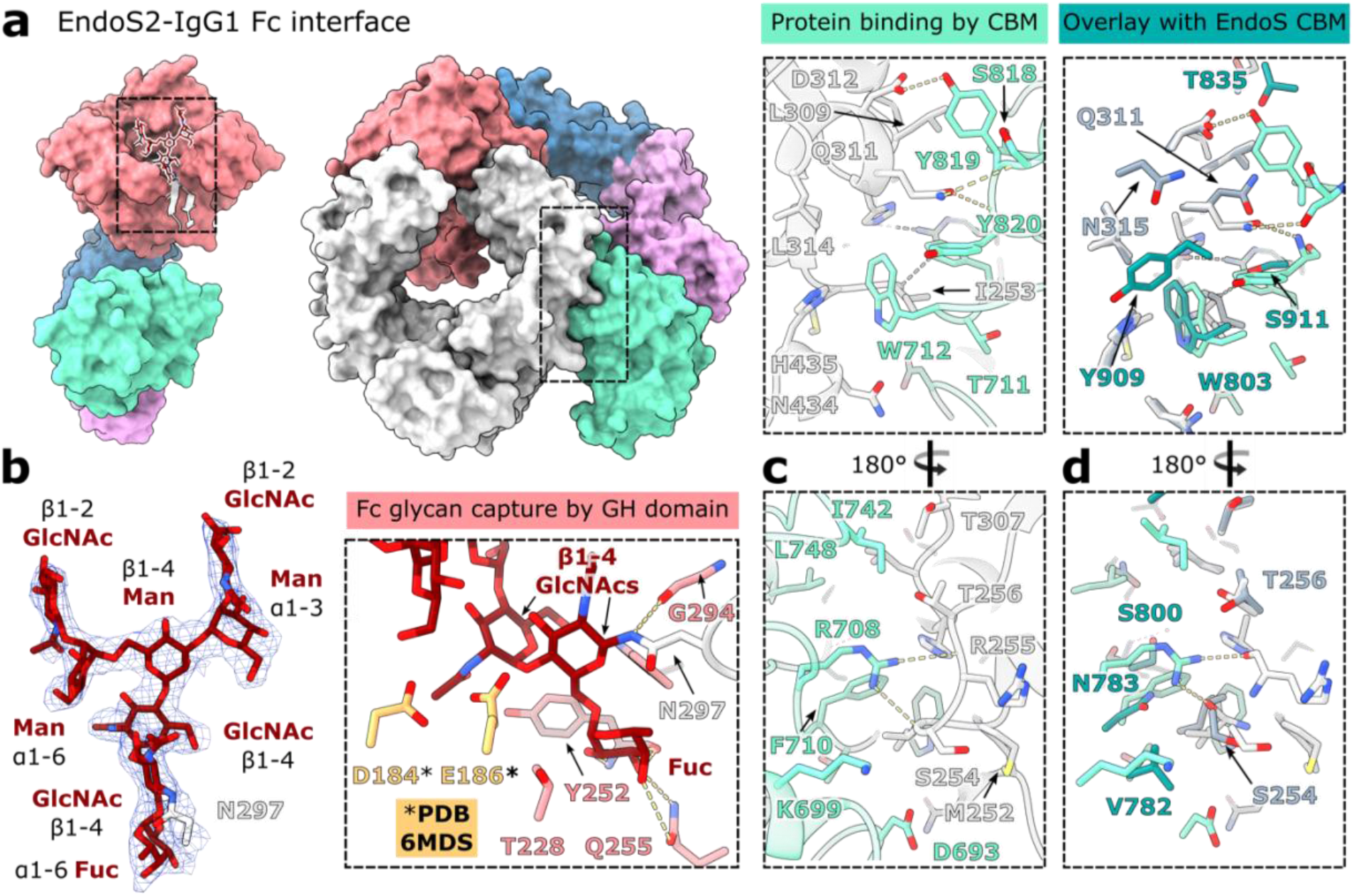
Binding interface of the EndoS2^D184A/E186L^-IgG1 Fc^L234C/E382A^ complex. **a** Overall structure of EndoS2^D184A/E186L^-IgG1 Fc^L234C/E382A^ complex, with interfacing areas highlighted. **b** N-linked glycan capture by the EndoS2 GH domain. Final 2*F*_obs_–*F*_calc_electron density map for the N-linked glycan and residue N297 is shown (weighted at 1.5 *σ*). Approximate positions of catalytic residues D184 and E186 (from an overlay with PDB 6MDS *via* the GH domain: aligned Cαs of residues 43*–*386) at the active site are indicated. **c** Fc protein surface binding by EndoS2 CBM. EndoS2 and IgG1 Fc are coloured as in Figure 1; residues involved in binding are shown as sticks and coloured by heteroatom. Hydrogen bonds are depicted as yellow dashes. **d** Overlay of EndoS2 CBM interface with EndoS CBM (PDB code 8A49). EndoS residues are coloured teal; IgG1 Fc complexed with EndoS is coloured dark grey. Residues involved in interface formation were identified using PDBePISA [71].

We observe atypical ϕ and Ψ torsion angles for the β1–4 linkage between the two core GlcNAcs of 5.9° and –121.2°, respectively, using C1–O–C(x)′–C(x–1)′ as the definition of Ψ. These values lie outside the range of average glycosidic linkages reported in crystal structures (–73.7° ± 8.4 for ϕ; 116.8° ± 15.6 for Ψ angles, calculated from 163 structures) [72]. In a catalytic context, the distortion of the torsion angles at the site of hydrolysis is consistent with promoting cleavage of the β1–4 linkage. Such glycan distortion is also observed within the EndoS-Fc complex, with equivalent ϕ and Ψ torsion angles of –58.6° and –121.2°, respectively [55]. In addition, several crystal structures of endoglycosidases in complex with their cleaved glycan substrate show the second GlcNAc adopting a higher-energy, skew-boat conformation, and so glycan distortion been suggested to comprise part of the catalytic cycle in such enzymes [11, 73, 74]. As observed for the structure of EndoS2 in complex with a complex-type glycan (PDB 6MDS [11]), the second GlcNAc is bound close to the catalytic dyad residues D184 and E186 (exchanged to alanine and leucine in our structure, respectively; Figure 2b), as indicated with an overlay of wild-type, apo EndoS2 (PDB 6MDS) via the GH domain. In this approximate position, E186 is oriented in close proximity to the distorted β1–4 linkage between the two GlcNAcs, facilitating nucleophilic attack and subsequent cleavage of the glycan.

The CBM interface in the EndoS2-IgG1 Fc complex is analysed in Figure 2c. Amino acid W712 in EndoS2 has been shown to be important for Fc binding, as substitution to alanine abolished hydrolytic activity towards IgG Fc bearing complex-type or oligomannose-type glycans [11]. The EndoS2-Fc structure reveals how W712 binds within a hydrophobic pocket at the Fc Cγ2-Cγ3 interface, formed by I253, H310, L314, N434 and H435. Thus, W712 makes similar interactions as W803 in EndoS, similarly shown to be essential for enzyme activity [53] and observed binding in the same cavity within the EndoS-Fc crystal structure [55] (Figure 2d). Another important amino acid in EndoS2 is Y820: Klontz *et al*. found that a serine exchange variant of this side chain severely reduced hydrolytic activity towards IgG bearing both complex-and oligomannose-type glycans [11]. This side chain makes not only a main chain contact to Fc residue Q311, but also forms stacking interactions with the Q311 side chain (Figure 2c). The equivalent amino acid in EndoS is a serine, similarly providing a main chain interaction with Q311 but lacking the hydrophobic interaction [55]. A further parallel between the two structures is observed with the side chain of EndoS2 Y819 that hydrogen bonds to the D312 side chain from the Fc, similar to the equivalent interactions of the EndoS T835 side chain [55].

Comparison of the EndoS2 and EndoS CBM interfaces with IgG1 Fc reveals some unique interactions formed by EndoS2, such as the side chain of R708 that hydrogen bonds to the Fc backbone (Figure 2c and 2d). PISA analysis [71] also predicts distinct Fc protein surface interfaces across the two enzymes: the EndoS2-IgG1 Fc interface has a surface area of 977.9 Å^2^ and a solvation free energy gain of –8.6 kcal/mol. Conversely, EndoS forms a 1323.5 Å^2^ interface with the Fc, with a –9.1 kcal/mol solvation free energy gain. A sequence alignment of EndoS and EndoS2 reveals that side chains interfacing with the Fc protein are not particularly conserved (Supplementary Figure 4). Moreover, aside from residue W712 (W803 in EndoS), other CBM residues found to be important for IgG Fc binding (R908 and E833 in EndoS [53]; Y820 in EndoS2 [11]) are not conserved (Supplementary Figure 4). These observations indicate distinct modes of IgG recognition by the two endoglycosidases.

Although the two enzymes bind IgG1 Fc in a similar way, a comparison of the EndoS2-Fc and EndoS-Fc crystal structures reveals differing angles of recognition between the two enzymes. Superposition of the two complexes with respect to the interfacing Fc Cγ2 domain (calculated by aligning Cαs of residues 237–340) shows how different GH domain and CBM orientations allow the enzymes to discover different surfaces on the Fc peptide (Figure 3a). This is accompanied with distinct Fc surface interfaces recognised by the two enzymes (Figure 3b). We additionally observe differential capture of the Fc by the two enzymes: superposition of EndoS2-and EndoS-bound Fc by their Cγ3 domains (alignment of Cαs, residues 341–444) reveals more “open” Cγ2 domain placements in the former (Figure 3c). However, an overlay of the interfacing Cγ2 (alignment of Cαs, residues 238–340) within the complexed Fcs and the equivalent domain from a wild-type IgG1 Fc structure (PDB 3AVE) shows that conformational changes only occur in the C′E loop, allowing capture of the Fc glycan in a “flipped-out” state (Figure 3b) [55]. A superposition of the GH domains from EndoS-and EndoS2-Fc complexes shows that the interfacing Fc Cγ2 domains are captured in different orientations (Supplementary Figure 5), which may reflect the altered Cγ2 domain placements observed here.

**Figure 3:**
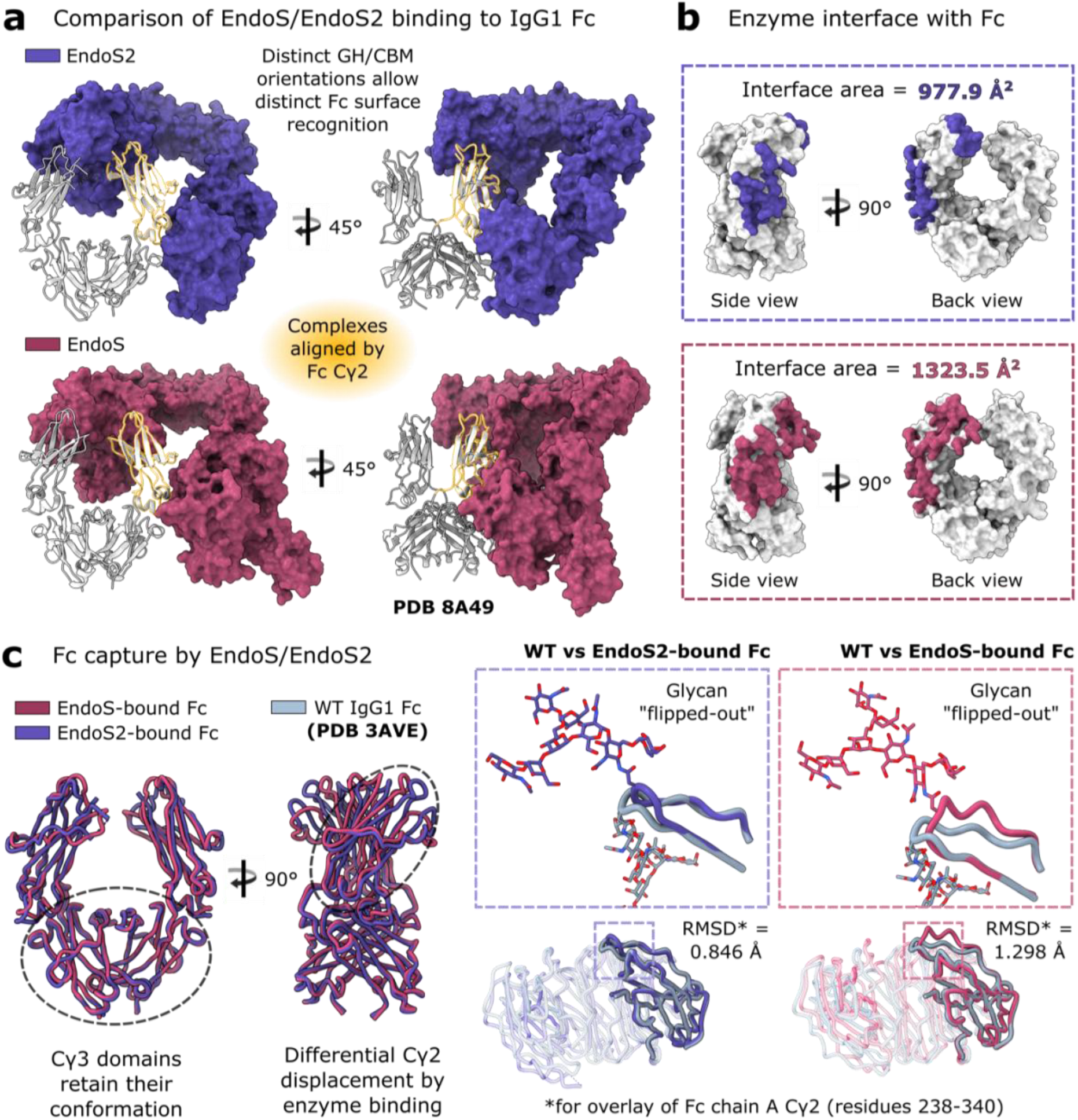
Comparison of EndoS and EndoS2 binding their IgG1 Fc substrate. **a** EndoS2-Fc and EndoS-Fc crystal structures, superimposed with respect to the interfacing Fc Cγ2 domain (Cαs for residues 237–340). Complexed EndoS (PDB 8A49; [55]) and EndoS2 are depicted in maroon and purple, respectively; complexed IgG1 Fc is depicted in silver. **b** Binding interfaces of each enzyme mapped onto IgG1 Fc, as calculated by PDBePISA [71]. **c** Comparison of Fc capture by EndoS and EndoS2. Superposition of EndoS-bound and EndoS2-bound Fc relative to the Cγ3 homodimer (Cαs for residues 341–444) reveals differential shifting of Cγ2 domains upon enzyme binding. Fc N-linked glycans (shown as sticks and coloured by heteroatom) are captured by both enzymes in a “flipped-out” conformation, occurring with shifts in the C′E loop which is covalently linked to the glycan (relative to a wild-type IgG1 Fc structure: PDB code 3AVE, depicted in dark grey). EndoS-bound and EndoS2-bound Fc are coloured maroon and purple, respectively.

The observed differences in EndoS and EndoS2 binding to IgG1 Fc, in distinct protein-protein interfaces and differing angles of recognition for the GH and CBM domains (Figure 3), provide a structural rationale for experiments with chimeric EndoS/EndoS2 enzymes, which showed that substitution of the EndoS2 GH onto an EndoS scaffold was not enough to confer EndoS2-like activity, but rather required substitution of both EndoS2 GH and CBM domains [11]. This chimeric enzyme was still not fully functional with respect to wild-type EndoS2 [11], thus the LRR–hIgG scaffold must also play a role in orienting the GH and CBM domains optimally. We envisage that our crystal structure can guide future efforts for synthesis of chimeric enzymes, which have application in engineering antibody glycosylation. Enzymes using EndoS/EndoS2 scaffolds are of particular interest due to their exquisite specificity for IgG antibodies: for example, a recent paper by Fan *et al*. shows how the glycosidase domains in EndoS/EndoS2 can be replaced with an α-L-fucosidase from *Lactobacillus casei* BL23 for significantly enhanced antibody defucosylation, when compared to the native fucosidase [75].

Taken together, our crystal structure of EndoS2 in complex with IgG1 Fc reveals the molecular basis behind its extensive substrate specificity. This structural knowledge will aid in the continued development of EndoS2, and chimeric EndoS/EndoS2-based enzymes, as tools for improved antibody glycoengineering and as biologics for use in immune modulation.

### Protein Data Bank accession number

Model coordinates and structure factors for the EndoS2^D184A/E186L^-IgG1 Fc^L234C/E382A^ crystal structure have been deposited in the Protein Data Bank with accession number 8Q5U.

## Acknowledgements

We are grateful to the beamline scientists on I03 at Diamond Light Source (DLS), and to Chris Holes for his support with the macromolecular crystallisation facility at the University of Southampton. We are also grateful to John Butler for his continued support and encouragement. This work was supported by DLS (MX29835) and the School of Biological Sciences, University of Southampton.

## CRediT authorship contribution statement

**Abigail Sudol:** Conceptualization, Investigation, Formal analysis, Writing - original draft, Writing - review & editing. **Ivo Tews:** Investigation, Formal analysis, Supervision, Writing - review & editing, Funding acquisition. **Max Crispin:** Conceptualization, Investigation, Formal analysis, Supervision, Writing - original draft, Writing - review & editing, Funding acquisition.

