## Supplementary Methods; Supplemental Table 1; Supplementary Figures 1-5 for "Bespoke conformation and antibody recognition distinguishes the streptococcal immune evasion factors EndoS and EndoS2"

*Cloning*. A gene fragment for IgG1 Fc, as encoded within a pFUSE-hIgG1-Fc vector, was synthesised by NBS Biologicals (plasmid purchased from Invivogen). An E382A exchange was included to inhibit preferential Fc self-crystallisation, as described previously [1]. The further L234C exchange was introduced in order to introduce an additional hinge disulphide (Supplementary Figure 1).

The enzyme EndoS2$_{38–843}$ was expressed with an additional C-terminal leucine (as described previously [2]) and a C-terminal linker and His-tag (sequence LEHHHHHH; Supplementary Figure 1); the amino acid exchange variant D184A/E186L was used to inactivate the enzyme (genome accession number ACI61688.1, cloned into a pET21a(+) vector by NBS Biologicals).

*Protein expression*. IgG1 Fc$^{L234C/E382A}$ was transiently expressed in Freestyle293F cells (ThermoFisher), using FreeStyle™ MAX Reagent (ThermoFisher), as described in the manufacturer's protocol. Cells were incubated at 37 °C, 8% CO2, shaking at 125 rpm (New Brunswick S41i incubator). Cell supernatants were harvested after seven days by centrifugation at 3100 × *g* for 30 minutes and filtered through a Nalgene™ Rapid-Flow™ Sterile Disposable Filter Unit (0.2 µm; ThermoFisher).

EndoS2$^{D184A/E186L}$ was expressed in BL21 (DE3)pLysS cells (ThermoFisher). Cells were grown in the presence of 100 µg/mL ampicillin and 34 µg/mL chloramphenicol in Terrific Broth (Melford), until reaching an $OD_{600}$ of 0.8, at which 1 mM IPTG was added. Cells were left to shake at 200 rpm (Innova 43 R incubator, New Brunswick Scientific) overnight at 25 °C. Cells were harvested by 20 minutes of centrifugation at 6220 × *g*.

*Protein purification*. The IgG1 Fc protein was purified by affinity purification with a HiTrap Protein A HP column (Cytiva). Eluted antibody was concentrated in a Vivaspin 20 Centrifgual Concentrator (MWCO 30 kDa; Sigma).

Harvested bacterial cell pellets were resuspended in PBS containing 2 µg/mL DNAse1 (Sigma) and a pinch of lysozyme (Sigma). Cells were homogenised with a glass homogeniser and broken apart using a cell disruptor (Constant Cell Disruption Systems). Remaining cell debris and cell membranes were removed by centrifugation at 3100 × *g* for 20 minutes, followed by centrifugation at 100,000 × *g* for 60 minutes. Supernatant was filtered through a Nalgene™

Rapid-Flow$^{TM}$ Sterile Disposable Filter Unit (0.2 µm; ThermoFisher) and subsequently passed over a 5 mL HisTrap HP column (Cytiva) for Ni affinity chromatography. EndoS2 was further purified with size exclusion chromatography using a Superdex 200 16/600 column (Cytiva) equilibrated in 10 mM HEPES, 150 mM NaCl, pH 8.0. Fractions corresponding to the main peak were pooled and concentratred in a Vivaspin 20 Centrifugal Concentrator (MWCO 50 kDa; Sigma).

*EndoS2-IgG1 Fc complex formation*. Protein concentrations of EndoS2$^{D184A/E186L}$ and IgG1 Fc$^{L234C/E382A}$ were calculated using a DS-11+ Spectrophotometer (DeNovix), using molecular weight and extinction coefficient values provided by the ProtParam tool [3]. EndoS2$^{D184A/E186L}$ and IgG1 Fc$^{L234C/E382A}$ were combined in a 2:1 molar ratio and the resulting complex applied to a Superdex 200 16/100 column (Cytiva) equilibrated in 10 mM HEPES, 150 mM NaCl, pH 8.0. Fractions corresponding to the main peak were pooled for crystallisation .

*Crystallisation*. The purified complex was subsequently exchanged into a buffer containing 50 mM HEPES, 150 mM KCl, pH 7.5, and concentrated to 10 mg/mL prior to crystallisation, using a Vivaspin 20 Centrifugal Concentrator (MWCO 30 kDa; Sigma). Crystal trays were set up using an Oryx4 robot (Douglas Instruments) and grown at 21 °C using sitting drop vapour diffusion in 0.1 M carboxylic acids, 0.1 M buffer system 3, pH 8.5, precipitant mix 4 (condition G12 in Morpheus crystallisation screen [4], Molecular Dimensions).

*Structure determination*. Crystals were flash-frozen in liquid nitrogen prior to data collection, which was carried out on beamline I03 at Diamond Light Source (Oxfordshire, UK) under a 100 K cryostream at a wavelength of 0.9763 Å. Diffraction images were processed using DIALS [5]. The complex crystallised in space group $P4_32_12$, with three copies of the EndoS-half Fc complex within the asymmetric unit. The structure was solved by molecular replacement with the program Molrep [6] within the ccp4i2 suite [7], using 3AVE and 6E58 search models for IgG1 Fc and EndoS2, respectively. The model was improved using successive rounds of manual model building and refinement, using Coot [8] and Refmac5 [9], respectively. Refinement was implemented using local non-crystallographic symmetry restraints, defined translation-libration-screw groups (Supplementary Table 1) and restraints generated from PDB-REDO [10]. Electron density maps were calculated using map sharpening in Refmac5. One EndoS2 copy was not well-resolved within the density and thus could not be fully built (Supplementary Figure 2), similar to what was observed with the recent crystal structure of EndoS-IgG1 Fc [1]. The Coot carbohydrate module [11] and Privateer [12] were used to build and validate N-linked glycan structure.

**a** IgG1 Fc$^{L234C/E382A}$ (residues 221-447)

DKTHTCPPCPAPE**C**LGGPSVFLFPPKPKDTLMISRTPEVTCVVVDVSHEDPEVKFNW
YVDGVEVHNAKTKPREEQYNSTYRVVSVLTVLHQDWLNGKEYKCKVSNKALPAPIE
KTISKAKGQPREPQVYTLPPSREEMTKNQVSLTCLVKGFYPSDIAVEW**A**SNGQPEN
NYKTTPPVLDSDGSFFLYSKLTVDKSRWQQGNVFSCSVMHEALHNHYTQKSLSLSP
GK

**b** EndoS2$^{D184A/E186L}$ (residues 38-843)

MEKTVQTGKTDQQVGAKLVQEIREGKRGPLYAGYFRTWHDRASTGIDGKQQHPEN
TMAEVPKEVDILFVFHDHTASDSPFWSELKDSYVHKLHQQGTALVQTIGVNELNGR
TGLSKDYPDTPEGNKALAAAIVKAFVTDRGVDGLDI**AIL**HEFTNKRTPEEDARALNV
FKEIAQLIGKNGSDKSKLLIMDTTLSVENNPIFKGIAEDLDYLLRQYYGSQGGEAEV
DTINSDWNQYQNYIDASQFMIGFSFFEESASKGNLWFDVNEYDPNNPEKGKDIEG
TRAKKYAEWQPSTGGLKAGIFSYAIDRDGVAHVPSTYKNRTSTNLQRHEVDNISHT
DYTVSRKLKTLMTEDKRYDVIDQKDIPDPALREQIIQQVGQYKGDLERYNKTLVLTG
DKIQNLKGLEKLSKLQKLELRQLSNVKEITPELLPESMKKDAELVMVGMTGLEKLNL
SGLNRQTLDGIDVNSITHLTSFDISHNSLDLSEKSEDRKLLMTLMEQVSNHQKITVK
NTAFENQKPKGYYPQTYDTKEGHYDVDNAEHDILTDFVFGTVTKRNTFIGDEEAFAI
YKEGAVDGRQYVSKDYTYEAFRKDYKGYKVHLTASNLGETVTSKVTATTDETYLVDV
SDGEKVVHHMKLNIGSGAIMMENLAKGAKVIGTSGDFEQAKKIFDGEKSDRFFTW
GQTNWIAFDLGEINLAKEWRLFNAETNTEIKTDSSLNVAKGRLQILKDTTIDLEKMD
IKNRKEYLSNDENWTDVAQMDDAKAIFNSKLSNVLSRYWRFCVDGGASSYYPQYT
ELQILGQRLSNDVANTLKDLLEHHHHHH

**Supplementary Figure 1: Constructs used for EndoS2$^{D184A/E186L}$-IgG1 $^{FcL234C/E382A}$ expression and purification**. **a** IgG1 Fc containing L234C and E382A exchanges. **b** Truncated EndoS2 (residues 38-843) containing D184A and E186L exchanges to abolish catalytic activity. Amino acid variations are highlighted in orange; C-terminal linker and His tag in EndoS2 construct is highlighted in blue.

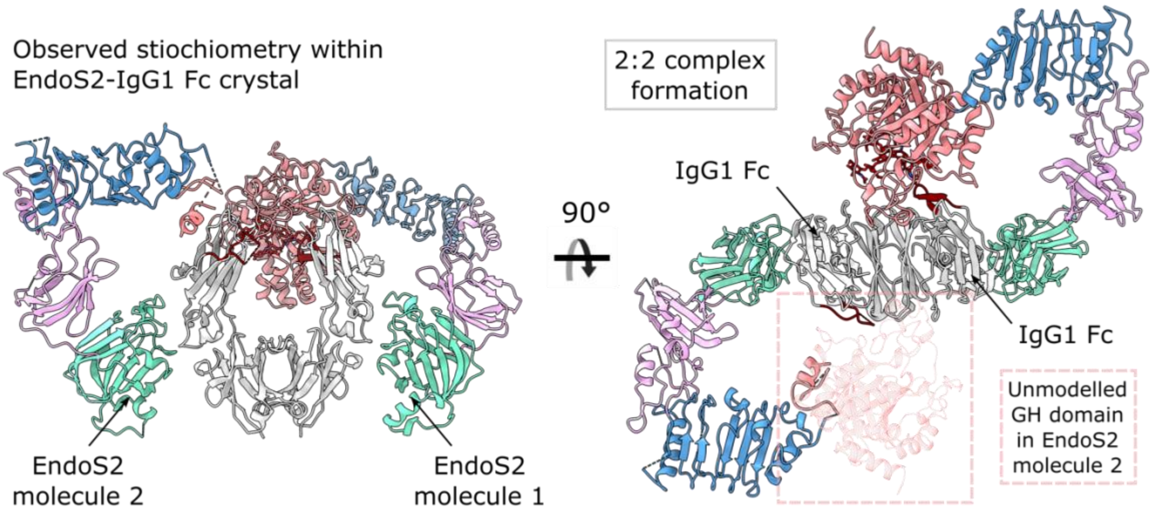


**Supplementary Figure 2: Stoichiometry within EndoS2$^{D184A/E186L}$-IgG1 Fc$^{L234C/E382A}$ crystal structure.** One EndoS2 molecule binds a single chain of the IgG1 Fc homodimer within the EndoS2$^{D184A/E186L}$-IgG1 Fc$^{L234C/E382A}$ crystal structure, resulting in a 2:2 stoichiometry as observed for the EndoS-IgG1 Fc complex [1]. EndoS2 and IgG1 Fc are coloured as in Figure 1.

**Supplementary Table 1: TLS parameters used in refinement of EndoS2$^{D184A/E186L}$-IgG1 Fc$^{L234C/E382A}$ crystal structure.**

| EndoS2$^{D184A/E186L}$ | IgG1 Fc$^{L234C/E382A}$ |
|---|---|
| TLS For Peptide chain D<br>RANGE ′D 45. ′ ′D 386. ′<br>RANGE ′D 387. ′ ′D 547. ′<br>RANGE ′D 548. ′ ′D 680. ′<br>RANGE ′D 681. ′ ′D 832. ′ | TLS For Peptide chain A<br>RANGE ′A 237. ′ ′A 340. ′<br>RANGE ′A 341. ′ ′A 444. ′ |
| TLS For Peptide chain E<br>RANGE ′E 264. ′ ′E 386. ′<br>RANGE ′E 387. ′ ′E 547. ′<br>RANGE ′E 548. ′ ′E 680. ′<br>RANGE ′E 681. ′ ′E 832. ′ | TLS For Peptide chain B<br>RANGE ′B 237. ′ ′B 340. ′<br>RANGE ′B 341. ′ ′B 444. ′ |
| TLS For Peptide chain F<br>RANGE ′F 46. ′ ′F 386. ′<br>RANGE ′F 387. ′ ′F 547. ′<br>RANGE ′F 548. ′ ′F 680. ′<br>RANGE ′F 681. ′ ′F 832. ′ | TLS For Peptide chain C<br>RANGE ′C 236. ′ ′A 340. ′<br>RANGE ′C 341. ′ ′C 443. ′ |

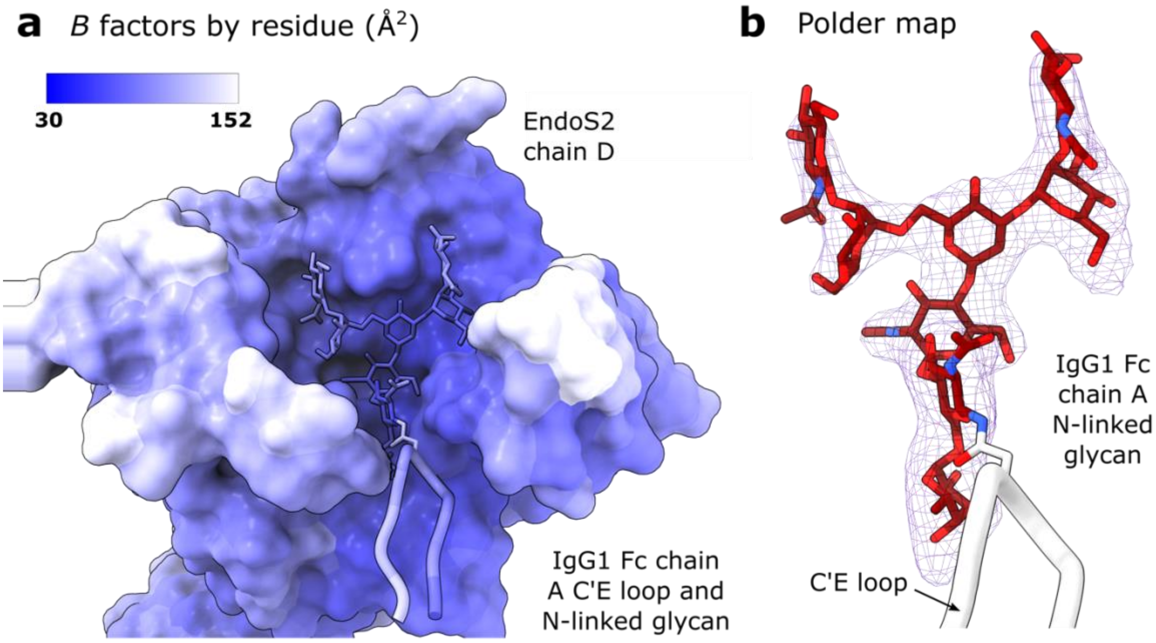


**Supplementary Figure 3: Validation of observed N-linked glycan conformation in EndoS2$^{D184A/E186L}$-IgG1 Fc$^{L234C/E382A}$ crystal structure**. **a** *B* factors per residue. **b** Polder map for the N-linked glycan modelled for chain A of IgG1 Fc$^{L234C/E382A}$, calculated (with default parameters) using the phenix.polder [13] in the PHENIX suite and shown at 3 $\sigma$.

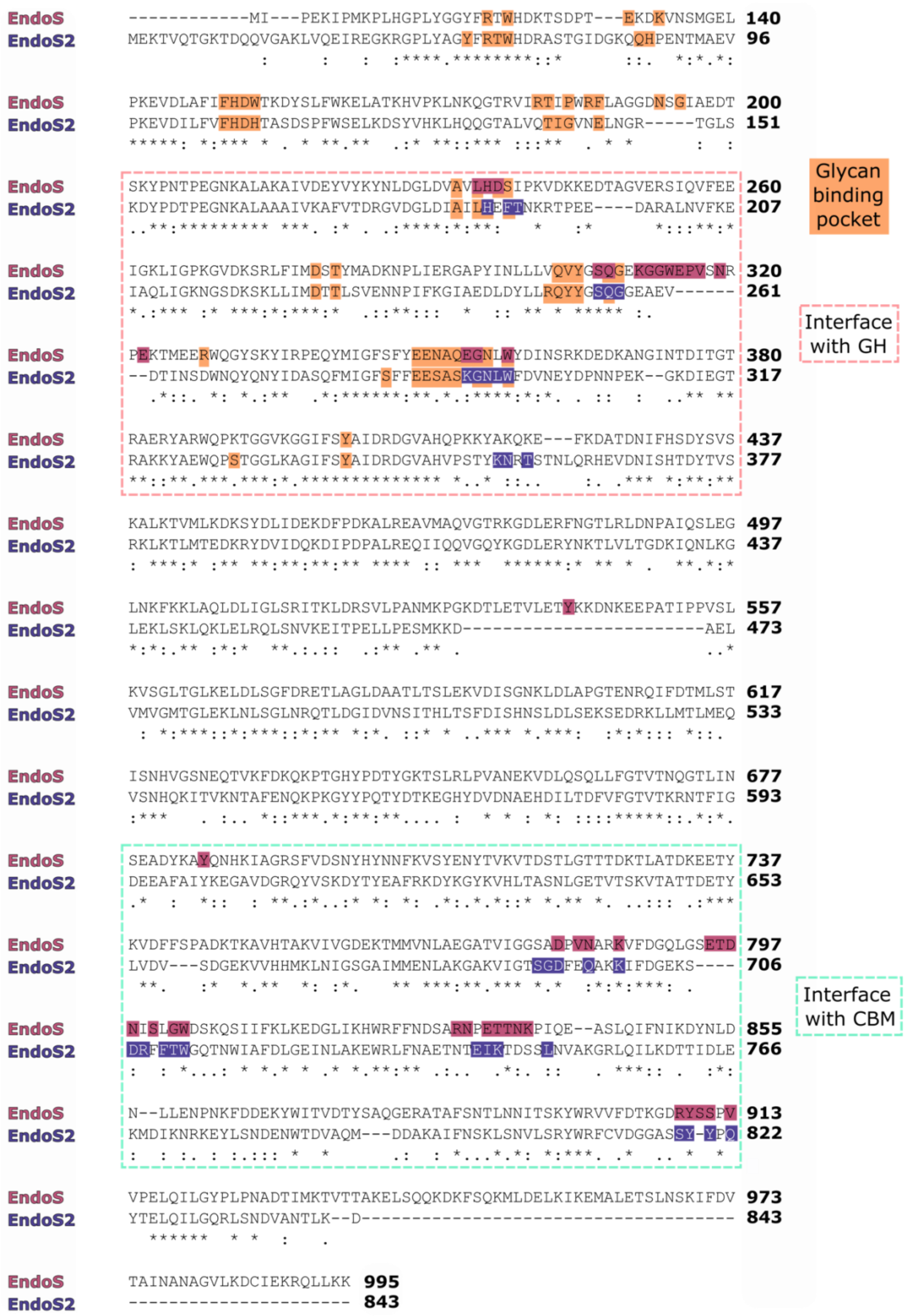


**Supplementary Figure 4: Multiple sequence alignment of EndoS2 and EndoS protein sequences, as generated by ClustalOmega [14]**. Sequences are shown as present in crystal structures of EndoS2$^{D184A/E186L}$-IgG1 Fc$^{L234C/E382A}$ (PDB 8Q5U) and EndoS-IgG1 Fc (PDB 8A49). Interfaces of EndoS2 and EndoS with the IgG1 Fc protein surface are coloured purple and maroon, respectively, with interfaces formed by the glycosyl hydrolase (GH) domain and carbohydrate-binding module (CBM) of each enzyme indicated with dashed lines. Interfaces of each protein with their glycan substrate are coloured in orange.

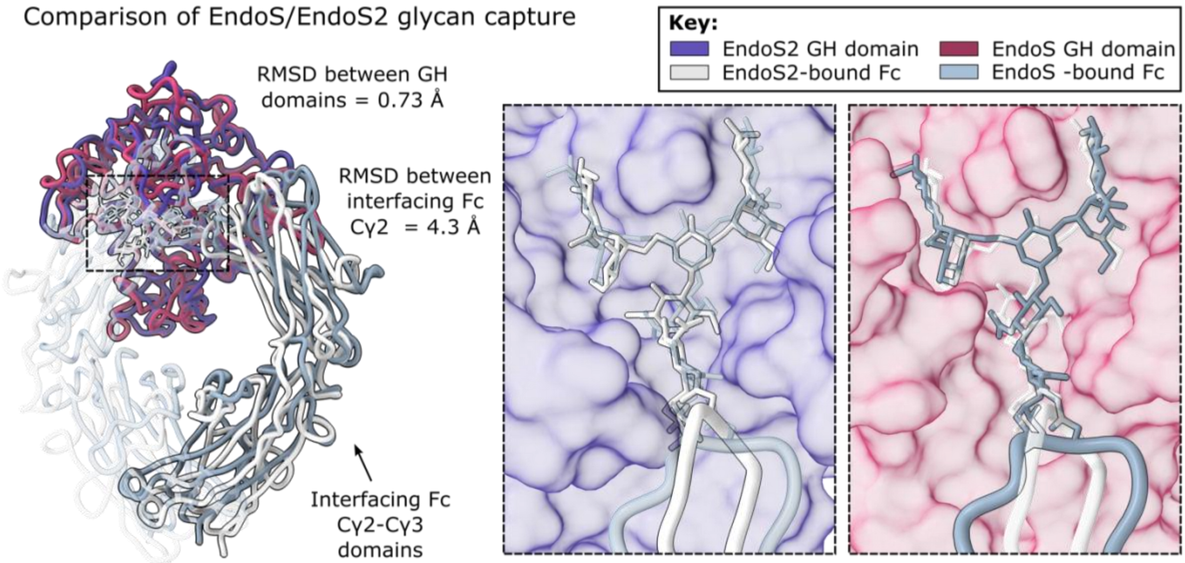


**Supplementary Figure 5: Superposition of GH domains from complexes of EndoS2 and EndoS (PDB 8A49) with IgG1 Fc.** Complexes were superimposed in ChimeraX [15] in relation to GH domains (amino acids 46–386 for EndoS2; amino acids 113–445 for EndoS), with a calculated RMSD of 0.73 Å between 273 pruned atom pairs. EndoS and EndoS2 GH domains are coloured as in Figure 3; EndoS- and EndoS2-bound IgG1 Fc are coloured dark grey and silver, respectively. Fc N-linked glycans are shown as sticks.
